# Targeting CDK12/CYCLIN K induces HIV gene activation and latency reversal which is mediated by P-TEFb

**DOI:** 10.64898/2026.02.10.705011

**Authors:** Praveenkumar Murugavelu, Heli Glebko, Jagrity Rani, Nili Tickotsky, Liron Levin, Alona Kuzmina, Ran Taube

## Abstract

The administration of antiretroviral therapy has successfully suppressed the replication of Human immunodeficiency virus and substantially inhibited viral infection. However, HIV still persists in long-lived infected cell reservoirs that are resistant to therapy and to immune clearance, therefore a barrier for elimination of infection. HIV gene expression is tightly regulated by the cellular transcription machinery, where low levels of host and viral transcription factors and epigenetic constrains maintain viral persistence. Here, we show that selective targeting of CDK12/CYCLIN K (CCNK) induces both HIV-specific and a global gene activation program, which is mediated by positive transcription elongation factor b (P-TEFb). Targeting CDK12/CCNK triggers a robust HIV gene expression, accompanied with latency reversal and synergistic reactivation when combined with latency-reversing agents. Mechanistically, CDK12/CCNK inhibition promotes the recruitment of RNA Polymerase II and CDK9 to the viral promoter, accompanied with elevated levels of histone activation makers. Targeting CDK12/ CCNK also exerts a global gene activation program, by releasing P-TEFb from 7SK snRNP, and reshaping the chromatin landscape at promoter-proximal loci and gene bodies. Together, these results uncover a previously unrecognized compensatory interplay between transcriptional kinases that rewires the cellular gene expression program with implications for HIV latency reversal.

**IMPORTANT:**

- Manuscripts submitted to Review Commons are peer reviewed in a journal-agnostic way.
- Upon transfer of the peer reviewed preprint to a journal, the referee reports will be available in full to the handling editor.
- The identity of the referees will NOT be communicated to the authors unless the reviewers choose to sign their report.
- The identity of the referee will be confidentially disclosed to any affiliate journals to which the manuscript is transferred.

**GUIDELINES:**

- For reviewers: https://www.reviewcommons.org/reviewers
- For authors: https://www.reviewcommons.org/authors

**CONTACT:** The Review Commons of**f**ice can be contacted directly at

## Introduction

HIV persistence remains a central obstacle for a complete cure for viral infection; while antiretroviral therapy (ART) effectively suppresses active viral replication, it fails to eliminate latently infected cells, which form cell-infected reservoirs mainly at resting memory CD4+ T cells that carry transcriptionally silent but replication-competent provirus. This reservoir persists for decades and therefore, the integrated intact provirus can rebound rapidly upon ART interruption, urging the requirement for a lifelong therapy^1,2^. A central mechanism for the induction of HIV latency is the low levels of host and viral transcription factors and the efficient block of transcription at the pause-release step, leading to overall suppression of HIV productive elongation (1, 2). As such deep understanding of metazoan gene transcription control is essential for revealing mechanisms of HIV gene expression regulation and establishment of viral latency. Indeed, cellular transcription program is a multi-step process, which is highly coordinated by the function of transcription factors and their kinase partners that jointly execute an accurate gene expression program in a highly dynamic environment (3, 4). Steps of the transcription cycle include initiation and recruitment of sequence-specific DNA-binding transcription factors (TF) to gene promoters, followed by binding of additional pioneering proteins and chromatin-modifying complexes that remodel the compact nucleosome and expose core promoter elements at transcription starting sites (TSS) for transcription. Key kinases play significant roles in promoting the transcription program. TFIIAH and Mediator associate with TF and Pol II, allowing Pol II to bind to the promoter and initiate nascent mRNA synthesis (4–9). CDK7 then phosphorylates serine residues at positions 5 and 7 (Ser5/7) of the C-terminal Domain (CTD) of Pol II (6, 10). While Ser5P attracts capping enzymes that act on the mRNA to disrupt interactions between Pol II and Mediator, to enhance promoter clearance and Pol II pausing; Ser7P primes the CTD for subsequent phosphorylation and transcription elongation events (11–13). DRB sensitivity-inducing factor (DSIF/SPT4/5) and the negative elongation factor (NELF) also assemble onto PIC to induce pausing of Pol II at +20-60 nucleotides from TSS (14–16). Only upon recruitment of the positive transcription elongation factor b (P-TEFb; CDK9; CYCLIN T) and Super Elongation Complex (SEC), the paused Pol II is released for efficient elongation. Pause-release of Pol II is catalyzed by CDK9 that phosphorylates SPT5 and NELF, as well as the Ser2-CTD (17, 18). Acting as a pivoted transcription factors within cells, the activity of P-TEFb is tightly regulated in cells. Active P-TEFb associates with super elongation complex (SEC), while the inactive complex resides within the 7SK snRNP and serves as a P-TEFb reservoir (19–22).

CDK12 and CDK13 also mediate Ser2 phosphorylation, enhancing transcription elongation (23–29). In cells, along with the role of CDK12 in transcription elongation, it also ensures proper expression of genes that are critical for stress response like DNA damage response (DDR), cell cycle progression and RNA processing (30, 31). Notably, CDK12 is distinguished from other transcriptional CDKs by its profound impact on genes involved in DNA repair pathways (32). Genome-wide analyses have shown that CDK12 occupancy overlaps with actively transcribed genes and enhancer regions, reinforcing its role as a master regulator of a specific subset of genes rather than exerting uniform control over the transcription program. Inactivation or loss of CDK12 functions have been linked to genomic instability, tumorigenesis, and altered cellular responses to DNA damage, making it not only a biological support in transcriptional regulation, but also a promising biomarker and therapeutic target for diverse human cancers)33-35(. These unique regulatory functions of CDK12 in transcription and DNA repair underline its importance in both normal cellular physiology and disease contexts (36, 37). Moreover, recent research confirms that inhibition of CDK12 stimulates the release of P-TEFb from 7SK snRNP, accompanied with induction of expression of host genes along with downregulation of DNA-damage response pathways (38, 39).

In this study, we show for the first time that specific targeting of CDK12/CCNK, a kinase/cyclin pair with key roles in transcription regulation of both cellular and HIV gene expression, induces global and HIV specific gene expression and promotes latency reversal through releasing P-TEFb (CDK9/CYCLIN T1) from its 7SK snRNP inhibitory complex. HIV infected cells were treated with SR-4835, an orally inhibitor of CDK12/CCNK that has optimal selectivity and few off-target effects when tested across a panel of 460 kinases. SR-4835 acts as a molecular glue that targets the interactions between the CDK12, CYCLIN K (CCNK) and CUL4-RBX1-DDB1 ubiquitin ligase complex, and facilitating targeted protein degradation of CCNK and overall depletion of CDK12 (40–42). SR-4835 is highly specific when compared to other currently tested CDK12 inhibitors like the covalent CDK12 kinase inhibitor, THZ531 (43). Additional molecular modeling studies using the X-ray structure of CDK12 predict that the binding mode of SR-4835 to be ATP is competitive, with SR-4835 interacting via hydrogen bonding with the hinge region of the kinase (Tyr-815, Met-816, and Asp-819). SR-4835 exhibits IC_50_ values for CDK12/13 at a range of 60nM, and 366nM for CDK13/CCNK, with overall with very low toxicity Importantly SR-4835 exhibits approximately 10-fold lower potency for CDK9 compared to CDK12 or CDK13, while displaying no inhibitory effect on other members of the transcriptional CDK, HIPK, and DYRK families (40–42).

Our results show that selective targeting of CDK12/CCNK by SR-4835 of HIV infected cells induces robust viral gene activation and latency reversal. Importantly, synergistic reactivation of latent HIV gene expression is detected upon inhibiting CDK12/CCNK and treating infected cells with standard Latency Reversing Agents (LRAs). Globally, treatment of cells with SR-4835 also induces a robust global gene activation program, which is mediated by the positive transcription elongation factor b (P-TEFb). Our results show that targeting CDK12/CCNK reshapes the epigenetic landscape around transcription starting sites (TSS) and gene bodies, inducing the release of P-TEFb from 7SKsnRNP. Additional transcriptomic profiling further demonstrates that CDK12/CCNK targeting upregulates P-TEFb target genes, specifically those that are short sized. Alongside, we document the suppression of DNA damage response pathways that are regulated by CDK12.

Overall, our findings suggest a novel interplay between cellular kinases that compensate each other and induces gene activation program, mediated by P-TEFb. Such gene activation program rewires the cellular gene expression program and has clinical implications for HIV latency reversal.

## Materials and Methods

### Cell lines and Culture Conditions

Jurkat, 2D10 and J-Lat 9.2 cells and GFP(+)-sorted HIV latent infected cells were cultured in RPMI 1640 (GIBCO) supplemented with 10% fetal bovine serum, penicillin-streptomycin and L-glutamine at 37C in humidified 5% CO2 atmosphere. See below culture conditions of primary CD4+ T cells.

### Antibodies

Antibodies were used for western blotting and cut and run: Anti-rabbit (Jackson Immunoresearch, Lot# 168738); anti-actin (Sigma, A5441, Lot # 030M4788); Anti-mouse(Jackson Immuno-research #115-035-062); Cyclin T1 (abcam, ab27963); Cyclin K (Cell Signaling #19472S); CDK12 (abcam-EPR29009-30 ab317746); Pol II CTD (Cell Signaling #2629S); Pol II CTD Ser2 (Cell Signaling #13499S); Pol II CTD Ser5 (Cell Signaling #13523S); CDK9 (Cell Signaling #2316S); H3K27ac (Cell Signaling #8173S).

### Isolation of primary CD4+ T cells

Human CD4+ T lymphocytes were isolated from healthy donors. PBMCs were isolated over a Ficoll gradient (Millipore) and were maintained at 2 × 10^6^ cells/mL overnight at 37°C. The following day, CD4+ T cells were isolated by negative selection with the RosetteSep Human CD4+ T-Cell Enrichment Cocktail (Stem cell Technologies), resulting in homogenous populations of CD4+ T cells. CD4+ T cells were cultured in complete RPMI media containing recombinant human IL-2 at 25 U/mL (Roche) to a final concentration of 10^6^ cells/mL and then stimulated using anti-CD3/CD28 Dynabeads (Invitrogen). Stimulated cells were counted, centrifuged for 5 min at 1,500 rpm and resuspended in fresh RPMI and IL-2 at a final concentration of 0.5 × 10^6^ cell/mL before transduction with high-titer HIV_GKO_ lentivirus. Transduced cells were cultured for an additional 48 h in complete RPMI media containing recombinant human IL2 and dynabeads at a ratio of 25 µL human beads per 10 million cells, before being analyzed by FACS for transduction levels.

### SR-4835 Treatment

2D10, J-Lat 9.2 and primary CD4+ cells were plated in 24-well flat bottom plates at a density of 1×10^5^ cells/ml supplemented RPMI 1640 media. Cells were treated with SR-4835 (at different concentrations) for 2 hrs and TNFα for 6 hrs. Cells were then washed with PBS and incubated for 48 hours. Analysis was performed by flow cytometry to measure GFP^+^ cells. For synergistic treatment, Cells were treated with 0.5uM SR-4835 for 2 hrs. Later washed with PBS and treated with 1uM LRAs (Prostatin, JQ1, SAHA) and Flavopiridol were treated to check whether it is CDK9 mediated activation.

### Cell Viability assay

Cell viability was measured using the XTT cell proliferation assay kit () according to the manufacturer’s instructions. Briefly 1 x 10^5^ cells were seeded per well in 24 well plate, treated with SR-4835 and incubated for 24 hours. XTT reagent was added, and absorbance was measured at 450nm after 3 to 4 hours using a microplate reader. Viability was calculated relative to untreated controls.

### Cell Apoptosis cell viability

Cell apoptosis was analyzed using Annexin V–pacific blue and Propidium Iodide (PI) staining. Cells were harvested, centrifuged at 1500 rpm for 5 min, washed once with cold Annexin V binding buffer, and resuspended in 100µL of binding buffer. Annexin V-FITC IgG (5µL) was added and samples were incubated for 15min at room temperature in the dark. Prior to acquisition, 100µL of binding buffer and PI (2.5µL) were added. Samples were analyzed immediately by flow cytometry.

### Protein immunoprecipitation

2D10 (5 x 10^6 cells) will be treated with 0.5 µM SR-4835 for 2 hours, followed by washing with PBS. After an additional 6 hours, the cells will be harvested and lysed using a lysis buffer containing RNase inhibitor and protease inhibitor. 10% of the total lysate will be collected as input control. The lysate will be incubated on ice for 10 minutes, after which antibodies against CDK9, HEXIM 1, AFF4, or a non-specific IgG control will be added. The samples will be incubated overnight at 4°C with rotation. The following day, magnetic Dynabeads (Protein A or G) will be added to the samples and incubated for 2 hours at 4°C. A magnet will then be used to collect the complexes, and the beads will be washed three times with a wash buffer. The immunoprecipitated complexes will be resuspended in a 5x reducing sample buffer and boiled for 5 minutes to denature the proteins. Finally, Western blot analysis will be performed to assess the protein interactions, including CDK9, HEXIM1, and AFF4, to determine the effect of CDK12 inhibition on P-TEFb complex components.

### RNA isolation and RNA sequencing

Total RNA will be isolated from the infected cells using Fenozol reagent according to the manufacturer’s instructions. Infected cells will be harvested and resuspended in Fenozol reagent. The samples will be mixed thoroughly to ensure complete lysis of the cells. After lysis, chloroform will be added to the samples, and they will be vortexed for 15 seconds. The samples will then be incubated at room temperature for 2-3 minutes before centrifugation at 12,000 x g for 15 minutes at 4°C. This step will facilitate phase separation. The upper aqueous phase containing the RNA will be carefully transferred to a new tube. An equal volume of isopropanol will be added to the aqueous phase, and the mixture will be incubated at −20°C for at least 30 minutes to precipitate the RNA. The samples will be centrifuged at 12,000 x g for 10 minutes at 4°C to pellet the RNA. The supernatant will be discarded, and the RNA pellet will be washed with 70% ethanol. The RNA pellet will be air-dried briefly and resuspended in RNase-free water or a suitable buffer. The quality and quantity of isolated RNA will be assessed using a spectrophotometer or an RNA quantification assay. cDNA synthesis will be done and followed by RNASeq libraries will be prepared using a library preparation kit.

### Chromatin Immunoprecipitation (ChIP-qPCR)

Treated and untreated cells were cross-linked with 1% formaldehyde for 25 min, and then cross-linking was stopped with glycine (137,5 mM; 5 min). Cells were washed twice with PBS, then lysed for 10 min on ice in 130μL sonication buffer (20 mM Tris [pH 7.8], 2 mM EDTA, 0.5% SDS, protease inhibitor cocktail (Roche). DNA was fragmented by sonication at the following settings: amplitude 40% for 30 cycles at 10s on/10 s off, and samples were centrifuged (15 min, 14,000 rpm, 4°C). The soluble chromatin fraction was collected and immunoprecipitated overnight at 4°C on a rotating wheel in IP buffer (0.5% Triton X-100, 2 mM EDTA, 20 mM Tris [pH 7.8], 150 mM NaCl, and 10% glycerol) with one of the indicated antibodies according to Manufactory instruction. Anti-Rabbit IgG was used as a background control. The next day, the IP material was incubated with 20 μL of protein G beads (Thermo 10003D) for 2 hrs. DNA was eluted with freshly prepared elution solution (1% SDS and 0.1 M NaHCO3) and heated at 65°C overnight to reverse-crosslink the samples. Precipitated DNA fragments were then extracted using a ChIP DNA clean and concentrator kit (ZYMO Research), and HIV DNA levels were quantified by qPCR with the primers specifically located on the LTR region of the HIV promoter. All signals were calculated as percent of input DNA.

### Cleavage Under Targets & Release Using Nuclease - CUT and RUN kit

CUT&RUN will be performed using a commercial kit from Epicypher. SR-4835 treated 2D10 cells will be analyzed in triplicate. After 24 hours, cells will be washed and coupled to Concanavalin-A coated beads, followed by overnight incubation in Digitonin (0.01%)-containing buffer with antibodies targeting non-specific IgG, H3K4me3, RNA Pol II (CTD, Ser 2, Ser 5) or CDK9. After incubation, cells will be washed with a permeabilization buffer and incubated with pAG-MNase for 15 minutes at room temperature. Additional washes will be performed, and CaCl₂ will be added to initiate chromatin cleavage at 4°C for 2 hours. The released DNA will be eluted, spiked with E. coli DNA, and used to construct barcoded libraries with the Epicypher library preparation kit. Finally, DNA will be quantified by Qubit. Libraries will be sequenced. Then sequences will be aligned to the combined human (Hg38), E. coli, and HIV reference genomes and normalized with IgG data. Enriched peaks for H3K4me3, RNA pol II (CTD, ser2 and ser5) or CDK9 will be identified using SEARCS. For visualization, bigwig files will be scaled and normalized using the E. coli spike-in DNA and will be visualized by Integrative Genomics Viewer (IGV).

### Data Availability

The analysis was carried out using the NeatSeq-Flow platform. Raw sequencing data in FASTQ format were trimmed by Trim Galore (version 0.4.5) (length = 25, q-25), and reads that were too short were discarded. FastQC, version 0.12.0, was used for read quality control. BWA Mapper (version 0.7.12, default parameters t = 20, B = 5) was used to align paired-end reads to reference human genome assembly hg38 and to the spike-in control (*Escherichia coli*) reference genome assembly. SAMtools was used to remove PCR duplicates and create coordinate-sorted BAM files. Normalization between samples and IgG was performed with a custom script that calculates read scaling factor based on spike-in DNA reads of each sample. The scaling factor was then used in the bamCoverage to normalize each sample according to its spike-in. Bigwig files were created by bamCoverage function from deepTools (version 1.5.91), and reads mapped to blacklisted areas designated by ENCODE were filtered. Reads that refer to off-chromosome locations were removed with BedClip (version 377) and sorted by Bedtools (version 2.30.0). Bigwig files were then created with bedGraphToBigWig by deepTools. ATF1 peaks were called by SEACR (version 1.3), the top 1% of enriched regions in target data were selected, and peaks 100 bp and less apart were merged. Peaks were viewed in Integrated Genomics Viewer (version 2.16.1) (and annotated with Homer (version 4.11. Peaks 500 bp upstream or downstream from the TSS were selected for further analyses. Matrices were generated with DeepTools createMatrix, and Heatmaps were plotted with deepTools plotHeatmap. The high-throughput sequencing data generated in this study have been deposited in the NCBI Gene Expression Omnibus (GEO) database. Both ChIP-seq and RNA-seq datasets are available under the accession number <u>GSE316746</u> and <u>GSE316689</u> respectively, allowing full access to the raw and processed data”.

### Statistical analysis

Data are presented as mean ± SD from three to four independent biological replicates. Statistical analysis was performed using two-way ANOVA in GraphPad Prism 7. Significance was defined as **** P<0.0001, **P ≤ 0.05; n.s., not significant.

## Results

### Targeting CDK12/CCNK with SR-4835 induces gene expression of latent HIV

To elucidate the role of CDK12/CCNK in regulation of gene transcription, we took advantage of the HIV gene expression program, which is heavily regulated by cellular machinery, primarily at steps of RNA pause-release; therefore, a model for studying gene transcription regulation. HIV infected cells were treated with a highly specific CDK12/CCNK inhibitor - SR-4835, which has been previously shown to selectively and high affinity target the CDK12 ATP binding site, promoting proteasomal degradation of CCNK only in the presence of CDK12 and the CUL4-RBX1-DDB1 E3 ligase complex)40(. SR-4835 was applied on two HIV infected Jurkat-derived CD4 T cells, which serve as models for studying HIV persistent. 2D10 cells are latently infected with an HIV-based lentiviral cassette, which expresses destabilized GFP reporter gene under the control of the HIV LTR and also encodes a mutant Tat (H13L) that suppresses gene transactivation, allowing HIV reactivation upon cell stimulation with standard LRAs or T-cell activation agents (29, 44). Jurkat (J)-Lat 9.2 cells harbor a replication-defective full-length HIV-1 provirus, where GFP is inserted instead of the NEF gene, enabling direct assessment of HIV proviral transcriptional activation from the integrated LTR upon T cell stimulation (45). Treatment of both cell types with increased concentrations of SR-4835 had no toxicity effects and did not affect cell viability; ensuring our ability to target CDK12 under the indicated conditions (**Fig. 1A**). We further confirmed by western blotting that indeed treating the T cells with SR-4835 depletes CDK12/CCNK protein levels (**Fig. 1B**). We then applied SR-4835 inhibitor to Jurkat T cells that serve as model for HIV latency. Treating 2D10 cells with SR-4835 induced a clear, dose-dependent increase in HIV-GFP expression, with reactivation levels rising substantially compared to untreated controls cells, or stimulation with TNFα (**Fig. 1C**). Similarly, in J-Lat 9.2 cells, where HIV is more difficult to reactivate, SR-4835 treatment led to a significant viral reactivation, as detected by increased levels of GFP expression (**Fig. 1D**). Finally, we also monitored HIV RNA levels by RT-qPCR in J-Lat 9.2 cells, confirming an increase upon treating cells with SR-4835 (**Fig. 1E**). These results demonstrate that CDK12/CCNK targeting by SR-4835 effectively reverses HIV latency.

**Figure 1.**
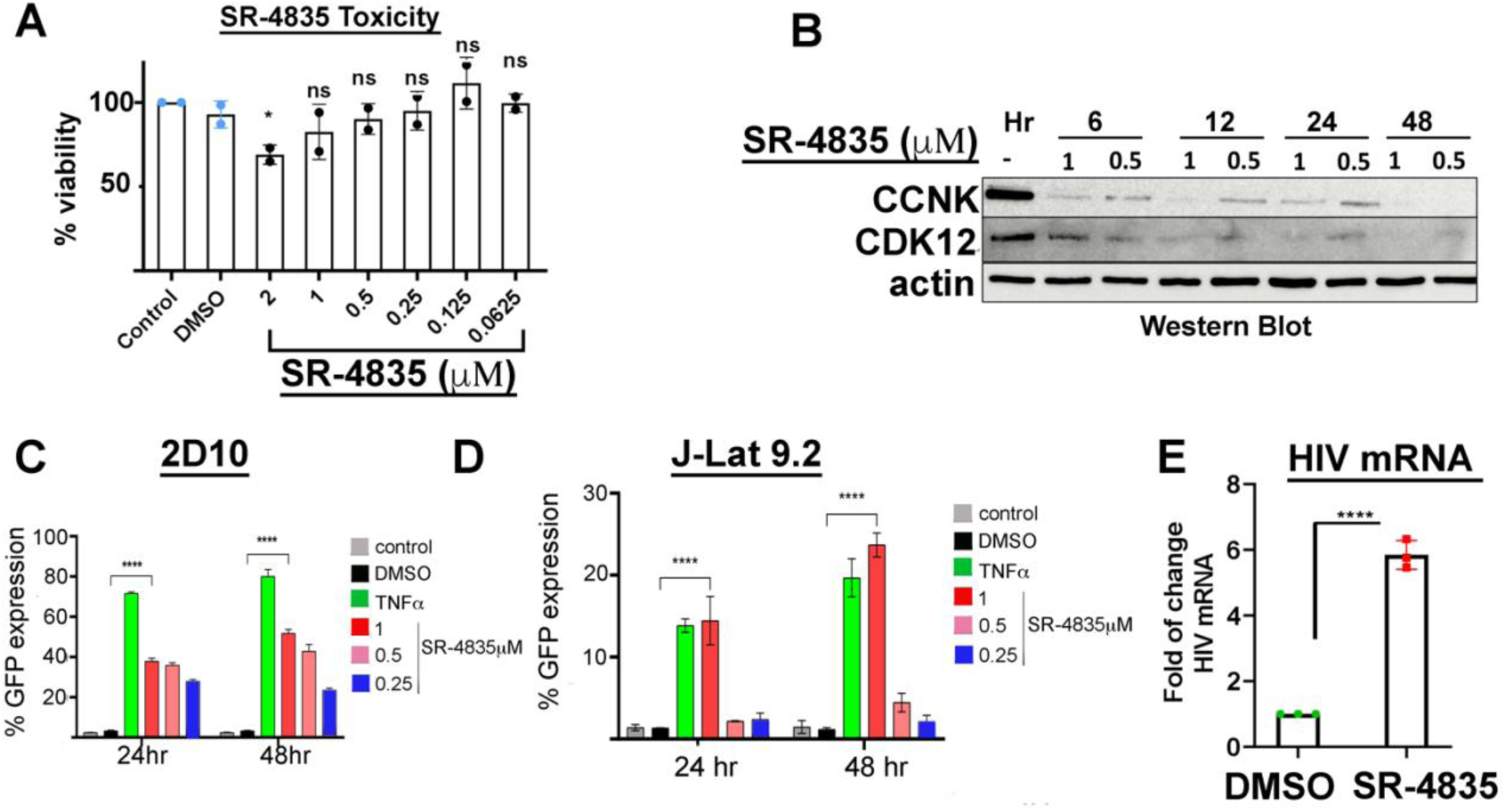
Targeting CDK12/CCNK reactivates HIV gene expression and reverses latency. **A.** Measurements of cell viability at 48 hours post SR-4835 treatment. XTT assays were performed on Jurkat T cells treated with increasing concentrations of SR-4835. Statistical significance was determined by two tailed t test. \*\**P* < 0.05 and ns: non-significant. **B. Levels of CDK12 and CCNK following SR**-**4835 treatment –** Jurkat T cells were harvested at the indicated time points (6;12;24;48 hours) post SR-4835 treatment, and subjected to Western Blot analysis with CDK12 and CCNK antibodies. **C-D**. 2D10 cells (**C**) and J-Lat9.2 (**D**) were treated with increasing concentrations of SR-4835. TNFα was used as a positive control for T cell activation and HIV reactivation. Cells were collected and monitored by FACS after 24 and 48 hours of treatment. Data represent mean ± SD from three biological replicates. Statistical significance was calculated using two-way ANOVA followed by Tukey’s multiple comparisons test (****P < 0.0001). **E**. HIV mRNA levels were detected in J-Lat 9.2 cells at 24 hours. of reactivation by qPCR and presented relative to GAPDH. Data represent mean ± SD from three independent experiments. Statistical significance was assessed using an unpaired two-tailed t-test. \*\*\*\**P* < 0.001.

### Targeting CDK12/CCNK with SR-4835 induces minimal cell apoptosis

Apoptosis and cellular stress may mimic HIV reactivation and latency-reversal (46). Therefore, it is critical to accurately identify whether SR-4835 treatment is truly reactivating HIV from latency, or is taking place following cell apoptosis. We therefore monitored apoptosis levels by FACS upon treating 2D10 cells with SR-4835, verifying that HIV reactivation indeed took place specifically in viable cells that did not go through apoptosis. Treated cells were analyzed by flow cytometry and gated only to live cells (Annexin V(-)/PI(-)), excluding apoptotic or necrotic perturbations in treated cells **(Fig. S1A**). Our observations also showed that treatment of cells at higher concentrations of SR-4835 led to minimal levels of apoptosis. Of note, HIV reactivation was monitored only in live gated cells **(Fig. S1B)**. This allowed us to assess the HIV activation independently of apoptosis, ensuring accurate measurement of HIV transcriptional activation in viable cells.

### Synergistic activation of latent HIV gene expression by combining CDK12/CCNK targeting and Latency Reversal Agents

We next tested effects of SR-4835-mediated reactivation of HIV latency upon combining treatment with currently applied Latency-Reversing Agents (LRAs). Cells were treated with SR-4835 and combined with LRA treatment, Prostratin (1μM), SAHA (1μM), or JQ1 (1μM), and I-BET (only for 2D10 cells, as other tested cells were not activated by this LRA) - each reactivating HIV latency via different pathways (47–51). We applied such a protocol on three experimental HIV latency models: 2D10, J-Lat 9.2 Jurkat T cell line, as well as Jurkat T cells infected with HIV_GKO_ and sorted for cells that carry latent HIV (GFP(-)/mKO2(+). HIV_GKO_ is a dual reporter full-length HIV lentivirus, where GFP is expressed under the control of the viral LTR promoter, while mKO2 reporter expression is drives by the general EF1α promoter (52). Cells were then treated with SR-4835 alone or in combination with the selected LRA, and HIV reactivation was monitored by FACS. In all three tested latency cell models, SR-4835 treatment induced a statistically significant increase in HIV - GFP(+) expression, indicating that it reverses HIV latency. Significantly, combined treatment of SR-4835 with LRAs, resulted in enhanced reactivation of latent HIV gene expression (**Fig.2**). In 2D10 the combined treatment of cells with SR-4835 and I-BET led to a significance increase (x6 fold). JQ1 and SAHA also exhibited a synergistic latency reverse reactivation when combined with SR-4835 (**Fig, 2A**). For J-Lat 9.2 cells, the combined treatment of SR-4835 with JQ1 led to the highest level of latency reversal, followed by combination treatments of SR-4835 with SAHA and Prostratin (**Fig. 2B**). Similarly, for Jurkat GFP (-)/mKO2(+) HIV infected T cells, we also detected an increase of HIV GFP expression upon combined treatment of LRA and SR-4835. The combination of SR-4835 with JQ1 or SAHA exhibited the highest increase in HIV-GFP expression compared to LRAs alone (**Fig. 2C**). These results suggest that SR-4835 has the potential to enhance LRA-mediated HIV gene activation and should be considered to be exploited in combination with LRAs to synergize HIV latency reversal.

**Figure 2.**
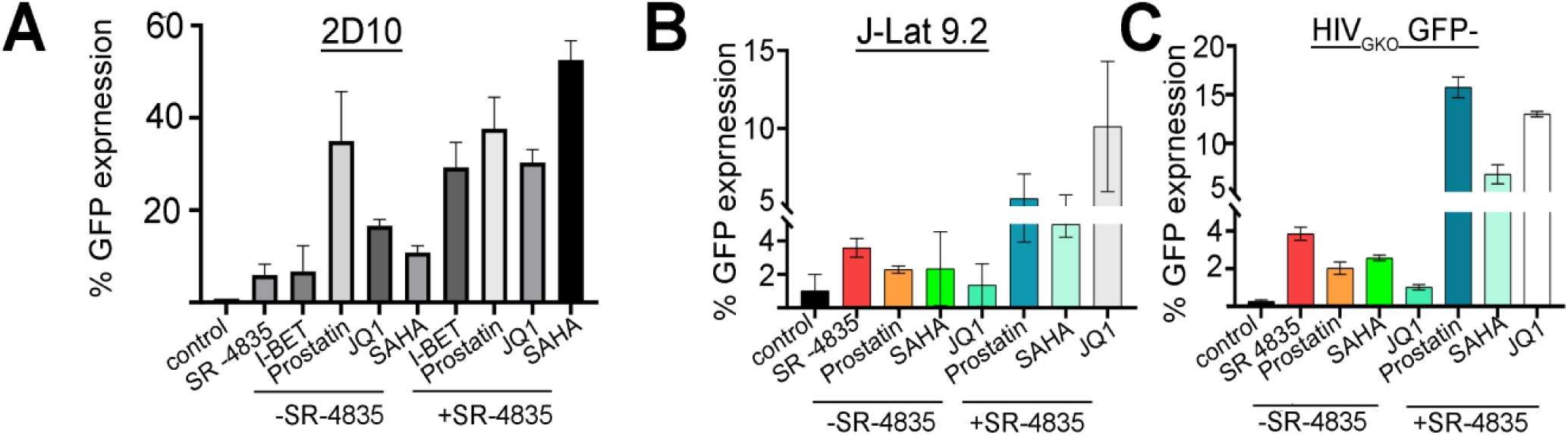
Targeting CDK12/CCNK synergizes reactivation effects of latency-reversing agents 2D10 (**A**); J-Lat 9.2 cells (**B**) and GFP-negative sorted HIV_GKO_ - infected Jurkat T cells (**C**), were treated with SR-4835 (0.5µM), Prostratin (1µM), SAHA (1µM), or JQ1 (1µM), either alone or in combination for 48 hours. 2D10 were also treated with I-BET inhibitor (1µM). HIV latency reversal was monitored by FACS based on HIV-GFP expression.

### Gene activation through targeting CDK12/CCNK is mediated by active CDK9

As P-TEFb/CD9 is key for HIV gene activation and RNA Pol pause-release, we sought to determine whether CDK9 kinase activity is required for the reactivation of HIV gene expression and latency reversal induced by CDK12/CCNK targeting by SR-4835. W treated cells carrying latent HIV infected latent cells [GFP(-);mKO2(+)] with SR-4835 (0.5µM) alone, or in combination with Flavopiridol (1µM), a potent and selective inhibitor of CDK9 kinase activity. 24 hours post treatment HIV latency reversal was monitored by FACS. Our analysis confirmed that upon targeting CDK12/CCNK with SR-4835, elevated expression of HIV-GFP was documented. However, SR-4835 mediated HIV latency reversal was efficiently suppressed (x7-fold) when cells were co-treated with Flavopiridol. Flavopiridol alone had minimal effects on HIV gene reactivation (**Fig. 3**). Flavopiridol inability to activate HIV gene expression, strengths the specificity of SR-4835 on CDK12/CCNK targeting. Overall, these results suggest that CDK9 kinase activity plays a role in SR-4835 induced latency reversal of proviral HIV (**Fig. 3**).

**Figure 3:**
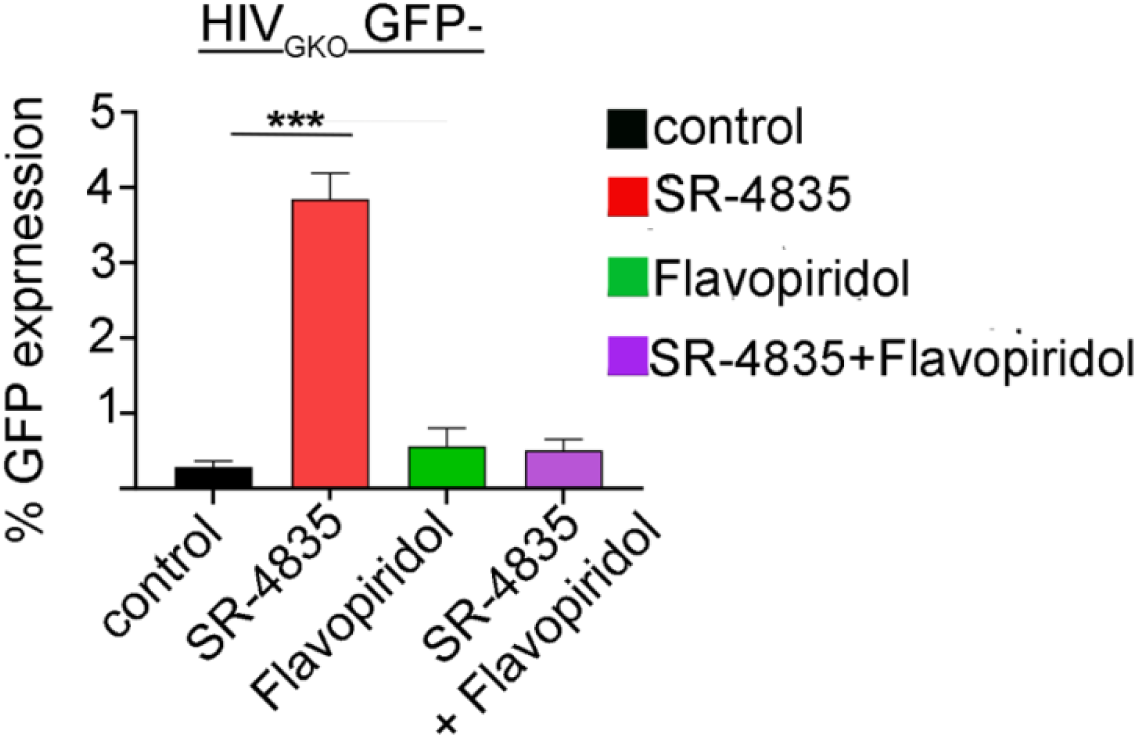
The kinase activity of CDK9 is required for HIV gene activation following CDK12/CCNK targeting by SR-4835 HIV_GKO_ infected Jurkat T cells were sorted for those where HIV is latent [(GFP(-);mKO2(+)] and then treated with SR-4835 (0.5µM) alone, or in combination with the CDK9 Flavopiridol (1µM) for 24 hours. HIV gene expression was monitored by measuring GFP expression by FACS. Data represent mean ± SD from three independent experiments. Statistical significance was assessed using an unpaired two-tailed t-test. \*\*\*\**P* < 0.001.

### Targeting CDK12/CCNK disrupts CDK9-HEXIM1 interactions and releases P-TEFb from 7SK snRNP

Upon establishing that targeting of CDK12/CCNK plays a key role in the reactivation of latent HIV through CDK9/P-TEFb, we aimed to investigate whether this effect was due to increased levels of CDK9 expression, or as a result of P-TEFb release from its inhibitory 7SK snRNP complex. To explore this, we performed co-immunoprecipitation (Co-IP)-CDK9 release experiments in Jurkat T cells treated with SR-4835. In non-treated cells, a significant fraction of CDK9 and CYCLIN T are sequestered in an inactive state within the 7SK snRNP complex, where HEXIM1 binds to CYCLIN T1 and inhibits CDK9 kinase activity)53(. This interaction prevents CDK9-mediated phosphorylation of Pol II at Ser2 of the CTD, thereby maintaining the transcription paused state of Pol II at TSS, including that of HIV (54). Our analysis included treatment of HIV infected cells with SR-4835, followed by IP of CDK9 from whole-cell lysates, and detection of associated HEXIM1 by Western Blotting (**Fig. 4**). Our results demonstrate that in SR-4835 treated cells, there was a marked decrease in HEXIM1 that was associated with CDK9 and CYCLIN T1, compared to untreated control cells (**Fig. 4A&B**). Importantly, total protein levels of CDK9 and HEXIM1 in the input lysates remained unchanged following SR-4835 treatment, confirming that the effect on HIV gene activation was due to the dissociation of CDK9/CYCLIN T1 from HEXIM1, rather than changes in expression levels. These results suggest that CDK12/CCNK targeting induces the release of CDK9 from the 7SK snRNP and activating gene expression.

**Figure 4.**
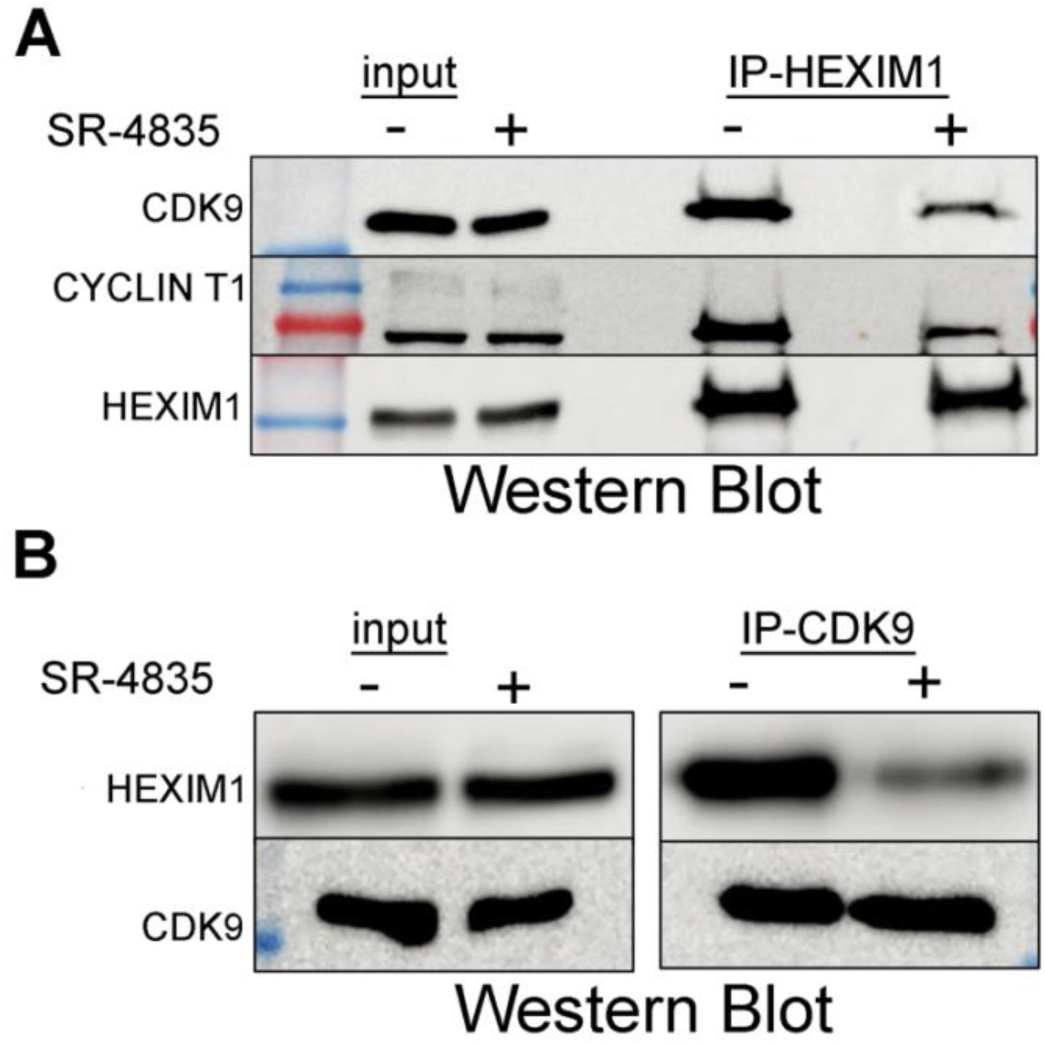
CDK12/CCNK targeting disrupts the association of CDK9 and CYCLIN T1 with HEXIM1. Jurkat T cells were treated with SR-4835 (0.5µM) or remained untreated for 24 hours, followed by immunoprecipitation with an anti-HEXIM1 IgG (**A**), or anti-CDK9 IgG (**B**). Co-IP fractions were analyzed by Western Blotting analysis for HEXIM1, CDK9 and CYCLIN T1.

### Genetic depletion of CDK12 or CCNK mimics SR-4835 treatment and activates HIV gene expression

To further confirm that CDK12/CCNK targeting and depletion activates HIV gene expression, we also took a genetic approach and depleted CDK12/CCNK expression levels in HIV infected cells (J-Lat 9.2) with siRNA specific for CCNK or CDK12. siRNA oligos (IDT) were electroporated into latently infected Jurkat T cells by electroporation (Neon NXT system), and efficient knockdown of CDK12/CCNK expression was confirmed by Western Blotting, showing a marked reduction in CCNK and CDK12 protein levels, compared to cells transfected with negative control siRNA (**Fig. 5A**). We further observed that upon depletion of CDK12/CCNK, HIV gene expression was induced, compared to non-targeted cells. Interestingly, combined knockdown of both CDK12/CCNK expression had no synergism effects on HIV gene expression, suggesting that loss of each component is sufficient to effectively activate HIV expression (**Fig. 5B**). Moreover, genetic depletion of CDK12/CCNK led to relatively minimal levels of HIV gene activation, implying that the knockdown was not complete, and minimal levels of these proteins are sufficient to activate HIV. These findings confirm that combined inhibition of CDK12 and CCNK, or depletion of each protein separately, induced reactivation of latent HIV infection.

**Figure 5.**
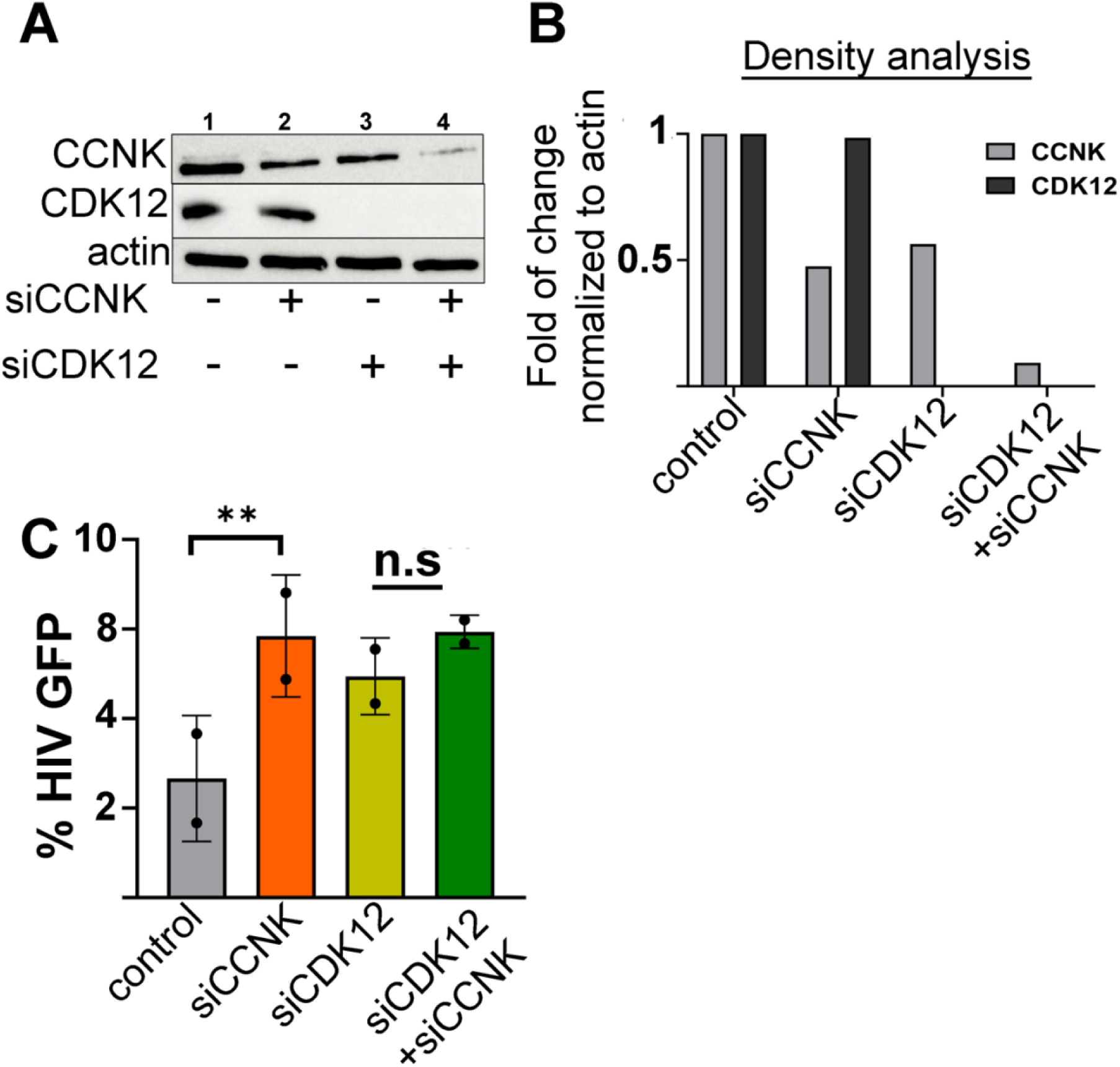
Genetic depletion of CDK12 or CCNK mimics SR-4835 treatment and activates HIV gene expression **A. Protein levels of CDK12 and CCNK following knockdown of CDK12/CCNK.** J-Lat 9.2 cells were electroporated with siRNAs specifically targeting CCNK (siCCNK), CDK12 (siCDK12), or both (lanes 2-4). Non-targeting siRNA served as control (lane 1). Protein levels of CCNK or CDK12 were monitored 48 hours post-electroporation by Western Blotting analysis. Monitoring actin levels served as a loading control. Densitometry analysis of bands is shown in panel **B**. **B. HIV latency reversal following CDK12 and CCNK protein depletion.** siRNA-electroporated J-Lat 9.2 cells were analyzed for HIV-GFP expression by FACS as a readout of HIV latency reversal.

### Targeting CDK12/CCNK reactivates HIV-transcription in primary CD4+ T cells

We next shifted our experiments and monitored the effects of SR-4835 on primary CD4+ T cells, which are the native target cells of the HIV reservoir. CD4+ T cells were isolated from healthy donors according to RostteSep^TM^ protocol and were then infected with the dual-fluorescent HIV_GKO_ lentivirus. At 48 hours post transduction, GFP (-); mKO2(+) cells were sorted and treated with SR-4835 to reactivate HIV from its latent state at low, non-toxic concentrations of SR-4835 (0.5µM). HIV reactivation in primary T cells was observed 24 hours post inhibitor treatment. FACS analysis of CD4 primary cells revealed a clear increase in HV-GFP expression upon SR-4835 treatment, compared to DMSO controls. These results demonstrated that the inhibition of CDK12/CCNK inhibition by SR-4835 reactivates HIV gene transcription in latently infected primary CD4+ cells (**Fig. 6A&B**).

**Figure 6.**
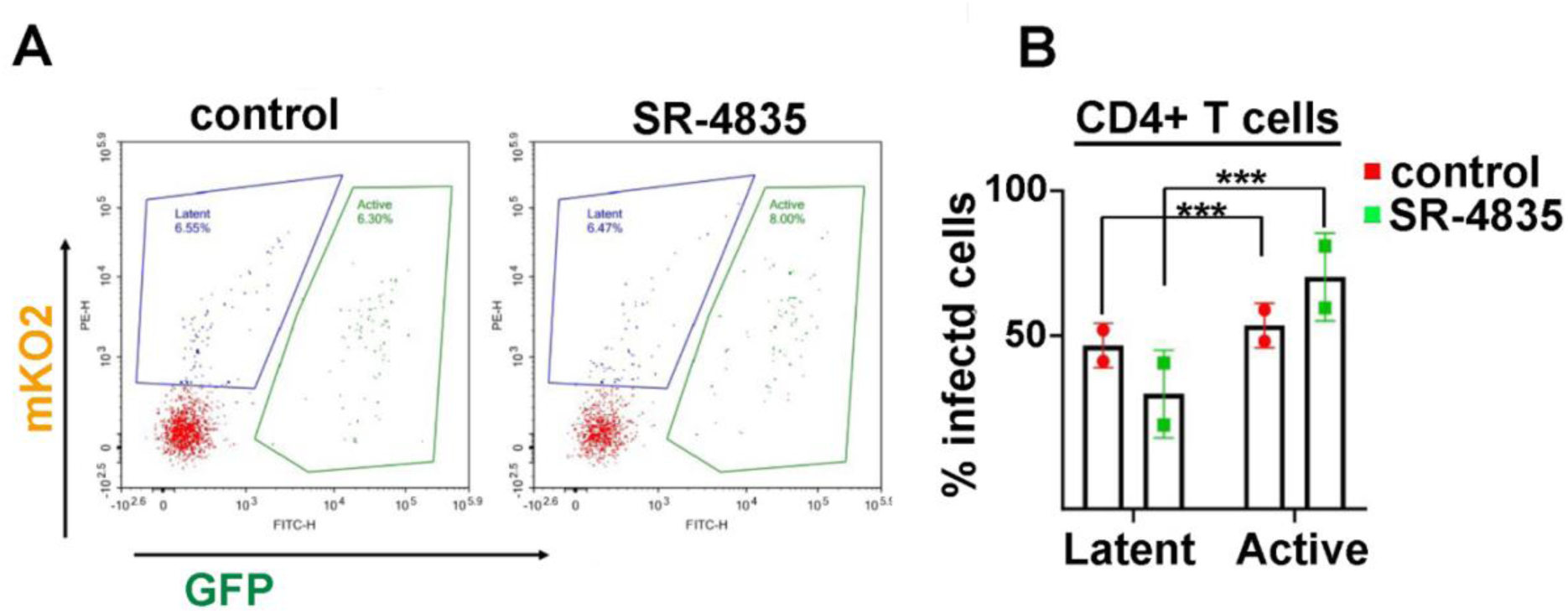
Targeting CDK12/CCNK with SR-4835 reverses HIV latency in primary CD4+ T cells. **A.** FACS analysis demonstrating that targeting CDK12/CCNK in latently primary CD4+ T cells which were transduced with HIV_GKO_ lentivirus and sorted for [GFP(-)/mKO2(+)] prior to SR-4835 treatment. Transduced cells were analyzed at 48 hours post transduction for HIV gene expression (GFP+/mKO2+). FACS dot plots presents HIV-GFP expression on the X-axis as a measurement of HIV gene expression and mKO2 expression on the Y-axis for transduction efficiencies. **B.** Quotative analysis of infection percentages (active HIV-GFP+) and latent (HIV-GFP-/mKO2+) normalized to infected cells (mKO2+) in SR-4835 treated cells compared to control non-treated cells.

### Targeting CDK12/CCNK induces the phosphorylation of Pol II and the recruitment of CDK9 and histone activation marks to the HIV promoter and to global gene promoters

Our results show that CDK9 is released from 7SK snRNP upon treatment of cells with SR-4835 and targeting CDK12/CCNK, potentially inducing transcriptional activation from the HIV promoter which is mediated by P-TEFb. To further confirm these results, we performed chromatin immunoprecipitation (ChIP) qPCR assays, monitoring levels of histone activation marks, as well as the occupancy of CDK9 and Pol II on the HIV promoter in Jurkat cells infected with HIV_GKO_. Cells were treated with SR-4835 for 24 hours, and chromatin was isolated and precipitated with antibodies targeting, H3K4me3 (epigenetic mark for active promoters), Pol II (total), CDK9, as well as for Ser2P and Ser5P that respectively mark initiation or elongation of transcription. Our data revealed that upon treatment of cells with SR-4835, higher enrichment was detected for H3K4me3 levels (x6-fold), indicating a transcriptionally an active environment at the viral promoter that follows CDK12/CCNK targeting (**Fig. 7A**). Similar results were obtained for CDK9 (x2.2-fold increase) upon SR-4835 treatment. These results are consistent with above observations of released P-TEFb from 7SK snRNP and recruitment of free complex to the HIV promoter to activate HIV gene expression (**Fig.4**). A similar increase upon SR-4835 treatment was also observed in levels of total Pol II (x4.7-fold); Ser2-P (x2 fold); and Ser5-P (x2 fold) (**Fig. 7A**). These results confirm that targeting CDK12/CCNK by SR-4835 induces HIV gene expression, accompanied with CDK9 recruitment and elevated levels of Pol II phosphorylation at Ser2 and Ser5, thereby probing Pol II for gene activation form the promoter HIV. To explore these results genome wide, we conducted CUT&RUN analysis in SR-4835 treated Jurkat T cells. Similarly, we observed elevated CDK9 occupancy at genome wide TSS and gene bodies (**Fig 7B**). Additional global analysis documented elevated levels at TSS and gene bodies of H3K4me3, H3K7ac histone activation marks, as well as Pol II, Ser2P and Ser5P, indicating enhanced transcriptional initiation and elongation activity.

**Figure 7.**
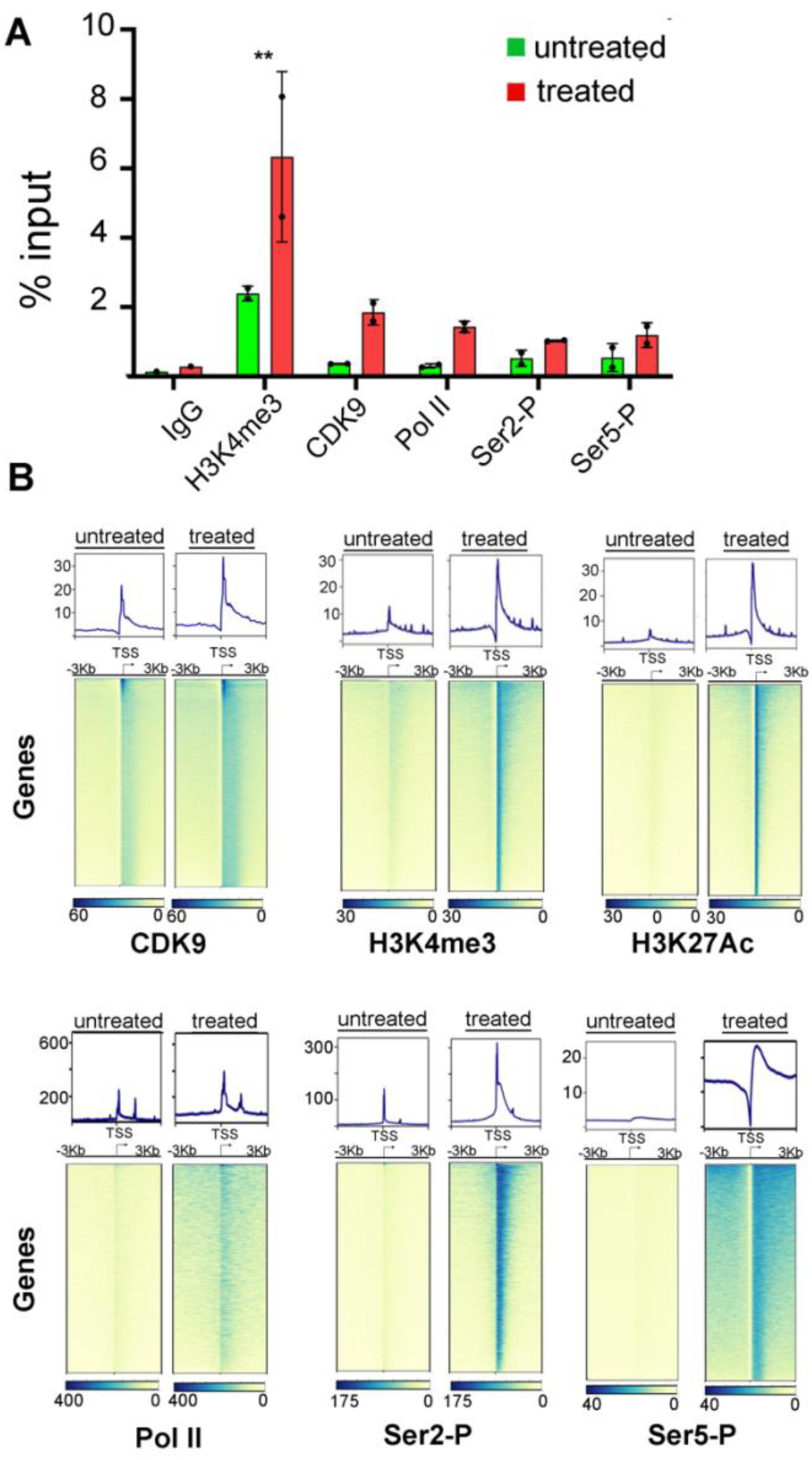
Targeting CDK12/CCNK induces HIV specific and global gene activation **A.** ChIP-qPCR analysis of Jurkat HIV transduced T cells treated with SR-4835 for 24 hours. ChIP qPCR was analyzed on ChIP material from SR-4835 treated, or untreated cells using the indicated antibodies. Data is presented as percentage of input relative to IgG as a negative control. Statistical significance is based on calculating mean ± SD from three independent experiments using One-way ANOVA. \*\**P* <0.01. **B.** CUT&RUN analysis shows global increase levels in gene promoters and gene bodies of CDK9 total RNA Pol II and well specific CTD Ser2-P and Ser5-P levels following SR-4835 treatment. Experiments were performed as described in the Methods based on Epicipher protocols.

### Targeting CDK12/CCNK alters global gene expression program

CDK12/CCNK play pivoted roles in regulating global metazoan gene expression program. To investigate the genome-wide transcriptome profile following SR-4835 inhibition, we performed RNA-Seq in Jurkat T cells infected with HIV. Our analysis identified, 30,226 DEGs (adjusted p<0.05) that upon SR-4835 treatment relative to the control untreated cells. Among these DEGs, there was a relative similar change between up and down regulated genes following SR-4835 treatment. We identified 1,448 (4.8%) upregulated genes and 2,059 (6.8%) down-regulated genes (**Fig. 8A**). Gene Ontology (GO) enrichment analysis identified significant enrichment of pathways associated with transcriptional regulation and activation, stress response and cell proliferation, while pathways related to DNA repair and replication were suppressed **(Fig. 8B)**. To visualize the expression patterns of the integrated HIV, we generated a heatmap of significantly altered viral genes following SR-4835 treatment, confirming increased expression of downstream HIV genes, specifically GFP, which is inserted downstream instead of the *NEF* gene (**Fig 8C**). IGV snap images confirmed the elevated reads that correspond to the HIV-GFP downstream loci. Cellularly, key proto-oncogenes such as MYC, FOS and JUN were also significantly upregulated upon Cdk12/CCNK targeting, while DNA damages response (DDR) pathways including genes like BRCA1, FANCD2, and FANCI were downregulated (38, 55, 56) (**Fig**. **8D**). These results confirm that CDK12/CCNK targeting suppresses DDR gene expression, while enhances transcriptional programs associated with cellular stress and cell proliferation. Interestingly, based on the analysis, we found that following treatment with SR-4835, shorter transcripts were significantly enriched in the upregulated genes relative to down regulated transcripts. In upregulated genes, those that were below 2.5Kb (<2.5kb) consisted 66.85% (968 genes), relative to 33.1% of the long transcripts (>2.5kb; 480 genes); while in the downregulated genes, there was no significance difference between the short or long transcripts 37.5% (774 genes) versus relative to long transcripts (62,5%=1286) (**Fig. 8E**).

**Figure 8.**
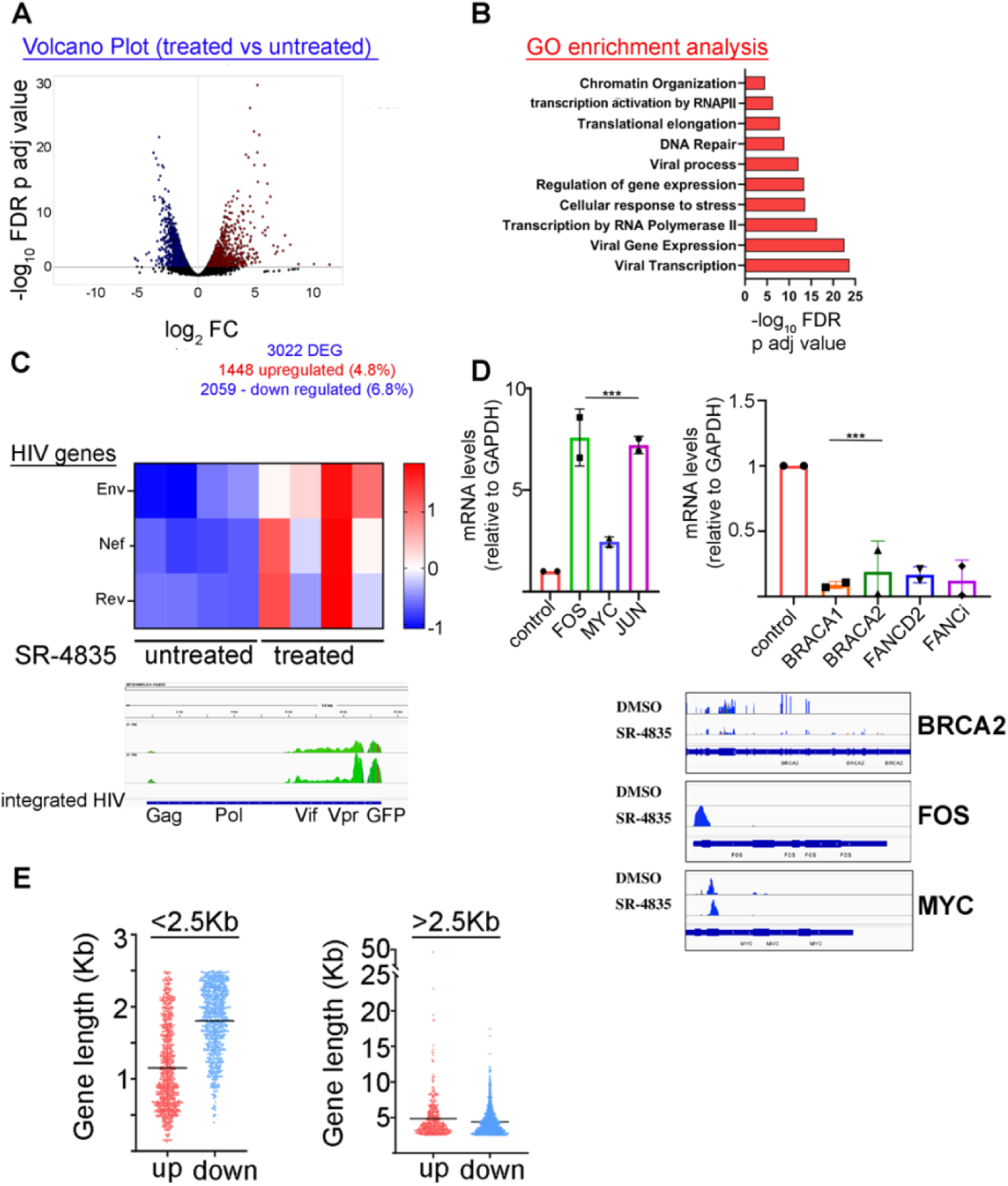
CDK12/CCNK targeting induces global changes of gene expression **A.** A volcano plot showing differentially expressed genes between SR-4835–treated and untreated cells. Red and blue dots represent significantly upregulated and downregulated genes, respectively. **B.** Gene Ontology enrichment analysis highlighting biological processes and pathways that are significantly modulated following targeting CDK12/CCNK by SR-4835. **C.** Heat map based on Z score, showing differential expression of HIV genes from the RNA-Seq transcriptomic dataset. GFP replaces the NEF downstream viral gene. **D.** DNA damage repair–related genes are downregulated upon SR-4835 treatment, while P-TEFb–target genes show increased RNA levels in SR-4835–treated cells. Control cells were untreated. Corresponding IGV peaks are also descried below. **E)** Analysis of gene size based on the RNA seq analysis in cells treated with SR-4835. Long transcrits are defined as >2.5Kb while shorter ones are <2.5Kb.

## DISCUSSION

In this work we employed a pharmacological approach to specifically target CDK12/CCNK and deplete their expression. Our results confirm that treatment of cells with a specific inhibitor - SR-4835 that acts as molecular glue, induces CCNK degradation and CDK12 depletion of expression with minimal toxicity effects and apoptosis. Moreover, targeting CDK12/CCNK expression by SR-4835 induces a dual and broad cellular gene expression cellular program, defined by marked downregulation of gene sets, but surprisingly induction of specific pathways and cellular genes. This gene activation program is manifested with an increase in genome-wide occupancy levels of Pol II, H3K4me3 and H3K27Ac activation histone marks around TSS as well as in gene bodies, which is accompanied with increased levels of CDK9, Ser5P and Ser2P of the Pol II CTD. Our results coincide with a recent unpublished work by Wang *et.a*l., showing a similar a transcriptomic profile upon targeting CDK12 (57). However, our study uses a more high affinity inhibitor for CDK12/CCNK, and further demonstrates that certain genes, primarily the short ones, are characterized with elevated activation marks and Pol II levels. Following CDK12/CCNK targeting, such specific gene activation program is driven by P-TEFb. Indeed, molecular characterization of the equilibrium of P-TEFb within cells, indicates that upon treating cells with SR-4835 and depletion of CDK12/CCNK, there is a shift of P-TEFb into its free configuration, releasing it from the inhibitory 7SK snRNP. Transcriptomic cellular profiling of genes that are differentially expressed upon CDK12/CCNK targeting, further show that there is a significant enrichment of cellular pathways that are associated with transcriptional activation, stress response and cell proliferation, while pathways related to DNA repair and replication, which are regulated by CDK12, are suppressed. Notably, key P-TEFb regulated proto-oncogenes such as MYC, FOS and JUN, as well as heat shock protein (HSP70) are significantly upregulated relative to control untreated cells, while multiple DNA damages response (DDR) genes including BRCA1, FANCD2, and FANCI are strongly downregulated. (38, 55, 56). Interestingly, based on the RNA-Seq analysis, upon treatment with SR-4835 there is a significant enrichment of short mRNA transcripts relative to longer genes, coinciding with results from Wang et.al. (57). These data highlight a key role of CDK12 in regulating an early transcriptional elongation checkpoint, where CDK12 targeting induces a premature termination and RNA decay that is disproportionally apparent in long genes; while, short genes show increased production of full-length transcripts. One of the notable short genes that were highly induced was MYC, known to induce potent pleiotropic effects and apoptotic cell death upon targeting CDK12 and CCNK. MYC gene expression following CDK12/CCNK inactivation may contribute to the specific killing of malignant cells, turning CDK12 as a potentially therapeutic target for treating human malignancies, where CDK12 is mutated or over-expressed such as HER2+ malignancies. Nevertheless, the therapeutic benefits of CDK12 targeting will require additional clinical investigation, given that those tumors readily develop resistance to multiple lines of other targeted therapies. The fact that P-TEFb is activated upon CDK12/CCNK targeting, where it reshapes a transcriptional response enabling survival of stressed cells, strengthen the therapeutic benefit of such CCDK12-CDK9 axis (58). Our results are consistent with recent reports on targeting CDK12/CCNK, demonstrating a broad gene activation effect primarily for short genes, including those encoding components of the AP-1 and NF-kB pathways in BRAF-mutated melanoma cells (39, 59) and genes driven by p53 and NF-κB (38). The novelty in our work is the understanding that targeting CDK12/CCNK drives P-TEFb to substitute CDK12 in activating a specific gene transcription program. This conclusion is further validated for specific upregulated genes, where levels of Pol II and Ser2P, are elevated upon targeting CDK12/CCNK.

Our study is also clinically significance, as it demonstrates for the first time that targeting CDK12/CCNK activates HIV gene expression and reverses latency. As targeting CDK12/CCNK drives the release of active P-TEFb and HIV is heavily dependent on P-TEFb-mediated Pol II pause-release and elongation of transcription, our results set HIV as an optimal model for studying the compensatory rewiring effects of CDK12/CCNK and CDK9/CCNT upon targeting CDK12/CCNK. Such effects are noted in several cell models for HIV latency, as well as in primary CD4+ T cells that carry a transcriptionally repressed provirus. Low doses of SR-4835 results in proviral gene activation, accompanied with elevated levels of Pol II, histone activation marks, as well as CDK9, Ser5P and Ser2P around the HIV promoter. Treating cells with SR-4835 combined with inhibiting the kinase activity of CDK9 with Flavopiridol, abolishes the ability of SR-4835 to reverse HIV latency. Moreover, synergistic effects on HIV gene activation and latency reversal were documented upon co-treating infected cells with specific LRAs and targeting CDK12/CCNK. LRAs are employed as part of a “Shock and Kill” strategy to target HIV infected cells where the provirus is transcriptionally silenced. However, while in-vitro LRAs are highly effective, they fail in the clinic (60, 61). We can speculate that a targeting approach of CCDK and CDK12, which is already employed for treating human malignancies, could be integrated in current tools to reactivate the infected reservoir - subjecting it to efficient killing by immune response and subsequent immune clearance. In addition to pharmacologic inhibition of CDK12/CCNK, we also genetically knockdown expression of CDK12/CCNK and monitored effects on HIV gene expression and latency reversal. Knockdown of CDK12 and CCNK, separately or together, results in only minimal HIV gene activation, implying that knockdown of the CDK12/CCNK is not sufficient for the activation patters detected with SR-4835 treatment. Overall, our study uncovers a key and novel interplay between CDK12 and CDK9, along with their cyclin partners. Upon targeting CDK12/CCNK, P-TEFb induces gene activation of set of specific genes, predominantly short genes, including HIV, while longer genes exhibit elongation defects that ultimately lead to premature termination. Such an interplay ensures a precise and coordinates gene expression program, where genes that are essential for stress response are induced.

## ACKNOWLEDGMENTS

We thank the people in the Taube lab for reading and revising the manuscript. This project is supported by the Bi-National Science Foundation (BSF, 2021273 to R.T.). Additional funding the Israel Science Foundation (ISF 1884/24 to R.T.). P.M, H. G, A.K, and J.R. and R.T designed and performed the experiments. N.T. and L.L. generated the data analysis and the metagenomics analysis. R.T. wrote the manuscript with input from all co-authors.

